# Matching dynamically varying forces with multi-motor-unit muscle models: A simulation study

**DOI:** 10.1101/2024.02.13.580042

**Authors:** T. Murtola, C. Richards

## Abstract

Human muscles exhibit great versatility, not only generating forces for demanding athleticism, but also for fine motor tasks. While standard musculoskeletal models may reproduce this versatility, they often lack multiple motor units (MUs) and rate-coded control. To investigate how these features affect a muscle’s ability to generate desired force profiles, we performed simulations with nine alternative MU pool models for two cases: 1) a tibialis anterior muscle generating an isometric trapezoidal force profile, and 2) a generic shoulder muscle generating force for a reaching movement whilst undergoing predetermined length changes. We implemented two control strategies, pure feedforward and combined feedforward-feedback, each parameterised using elementary tasks. The results suggest that the characteristics of MU pools have relatively little impact on the pools’ overall ability to match forces across all tasks, although performances for individual tasks varied. Feedback improved performance for nearly all MU pools and tasks, but the physiologically more relevant MU pool types were more responsive to feedback particularly during reaching. While all MU pool models performed well in the conditions tested, we highlight the need to consider the functional characteristics of the control of rate-coded MU pools given the vast repertoire of dynamic tasks performed by muscles.

## 1 Introduction

Vertebrate muscle is known for its mechanical versatility allowing it to perform athletic tasks of high speed, power, and endurance, but also delicate tasks requiring great finesse. How this versatility is achieved *in vivo* is not fully understood, although many structural and physiological characteristics affecting force production have been identified [1, 2]. One key factor affecting the control of muscle contraction is the compartmentalisation of a single muscle into multiple motor units (MUs), each comprising a motor neuron and the muscle fibres it innervates, and the recruitment of MUs in this pool a manner depending on task demands [3, 4]. In vertebrates, MU pool recruitment is extremely difficult to study *in vivo*, although recent advances in high-density surface electromyography (EMG) and blind source separation have enabled determining the recruitment patterns of multiple MUs during voluntary muscle contractions in humans (e.g. [5, 6, 7, 8, 9]). However, due to the technical challenges of EMG and other electrophysiological methods (see e.g. [10]), much of our current understanding of recruitment phenomena remains based on a small fraction of MUs in a pool [11], a limited number of muscles producing reliable data [11], and results that can be difficult to compare across studies [10]. Together with the common reliance of isometric tasks (e.g. [5, 12, 8, 6, 9]), these challenges limit our ability to infer the functional consequences of specific recruitment strategies across the range of contraction conditions in daily activities.

Computational MU pool models enable studying the behaviour of multi-MU muscles with more precise control and greater flexibility than experimental approaches. Existing models range from computationally efficient phenomenological models, most based on the model by Fuglevand et al. [13] (see e.g. [14]), to biophysical models considering muscle contraction at multiple scales from motor neuron membrane dynamics and cross-bridge cycling within muscle fibres, to continuum models capable of representing 3D deformations of the muscle [2]. Although biophysical models offer high physiological fidelity, they are computationally complex and have large parameter spaces, making them unsuitable for investigations requiring extensive parameter sweeps or a very high number of simulations. Hence, phenomenlogical models have been used to study phenomena such as onion-skin firing patterns (i.e. later recruited Mus reaching lower firing rates than earlier recruited ones) [15], variability in force production [16, 17], time-dependent changes and fatigue [18, 19, 20], and contraction energetics [21]. The common features of these MU pool models include threshold-based recruitment according to the size principle [22] and an explicit rate coding function (i.e. the relationship between the neural input driving MU recruitment and the motor neuron firing rate) which can vary from simple linear [13] to piece-wise linear [17] to mixed linear-exponential [23]. An intermediate option between phenomenological and full motor neuron membrane models is offered by leaky integrate-and-fire models, where an excitation impulse is triggered when the motor neuron membrane voltage reaches a threshold [12, 24]. Consequently, firing rate in these models is an emergent characteristic, rather than explicitly modelled, while the order of MU recruitment remains threshold-based. Regardless of the modelling approach taken, current MU pool models focus on offering insight into muscle contraction for a given neural input without explicitly addressing the control of the input during functional tasks. Hence, they cannot be used to unravel the links between MU recruitment strategies and the resulting movements alone.

Musculoskeletal (MSK) models are used widely to understand human movement and its control across different ages and health conditions (e.g. [25, 26, 27]). MSK models with multiple MUs have been used in to study phenomena such as sinusoidal single-joint movements [28, 29], MU recruitment based on metabolic cost [30, 31, 32], and standing balance [33], although most models rely on only two parallel MUs [30, 31, 32, 33] and are used for functionally simple tasks. Multi-MU MSK models utilising EMG-driven input (e.g. [34, 35, 24]) can achieve better physiological realism and functional versatility, but they cannot be used truly predictively and their usability is limited by the technical challenges of EMG acquisition mentioned above. While multi-MU MSK models exist, most models rely on representing muscles using a single Hill-type actuator for each direction of action, controlled via a continuous, amplitude-coded excitation or activation signal [36]. While approaches utilising one or a few MUs are computationally and parametrically convenient, they have limited power to address the functional consequences of multi-MU properties and rate coding.

Integrating existing models of full MU pools into MSK models carrying out a variety tasks is challenging because MU pool models have mostly been tested with isometric tasks requiring constant or slowly varying force production, such as constant excitation [13], sustained contraction [18], trapezoidal force production (i.e. ramp up - hold - ramp down) [12, 24], and step-wise force-level increase tasks [23]. Although MU pool models have been combined with tendon dynamics and Hill-type force-velocity-length (FVL) characteristics [21, 17, 24], the long, constant excitation [17] or target force [21, 24] conditions studied result in the MUs operating isometrically for most of the task. Full MU pool models have been successfully used to control a ballistic movement of a single joint [29] but without accounting the effect of muscle FVL characteristics on force production. Compared to the tasks used to study MU pool models, daily movements require a wider range of contraction conditions, including notable changes in muscle length and speed during contraction as well as changes in force production that can occur at very short time scales.

An additional challenge to using multi-MU muscles in MSK simulations arises from the practicalities of controlling MU pools during dynamic tasks which are largely unexplored. MU pool models with feedback control have been able match constant reference force levels [23, 21, 19, 18, 37] and replicate overshoot features commonly produced by humans [21], but the focus has remained on long, sustained force production tasks or intermittent contractions at constant force, with little attention given to the transient behaviour before the system reaches steady state. However, the time scale of many daily movements requires muscles to operate outside steady-state conditions for the majority of the movement; hence, control approaches accounting solely for the steady-state characteristics of MU pools are not guaranteed to work. An alternative approach to controlling a full MU pool is a hierarchical optimisation of the timings of individual MU twitch forces [28, 29], but it is computationally heavy for large MU pools and, to our knowledge, has not been proven to work when the significant nonlinear effects arising from muscle FVL properties and the summation of twitches are included. Two-MU muscles have been used to model cycle pedalling, controlled via optimisation of metabolic cost [31, 32], and standing balance, controlled using a model of proprioceptive reflex loops [33]. While these studies provide valuable insight into the role of fast and slow MUs, it is not clear how well the control approaches would translate to realistically sized MU pools of hundreds of MUs.

In the present work, we investigate whether the differences in the structure and properties of various MU pool models affect their ability to generate a desired force profile precisely, and to what extent any differences observed are task dependent. To answer these questions, we use computational modelling to compare different MU pools in two cases: (i) a tibialis anterior (TA) muscle required to match isometric trapezoidal forces, which can be compared with experimental data [5], and (ii) a generic (horizontal) flexor of the shoulder (SF) required to match the forces from simulated reaching movements [38], capturing realistic time scales and non-isometric muscle conditions for MSK modelling. To achieve a high level of interpretability, we investigate the MU pools’ ability to match force profiles in time-based simulations while undergoing pre-determined length changes, rather than embedding the multi-MU muscles within MSK models where nonlinear system dynamics confound outcomes. In the present work, we focus on how the recruitment and rate coding properties of MU pools impact force control (differences fibre contractile properties is beyond the current scope). With an eye towards eventual integration of MU pool models with MSK models, we develop and test two control strategies of MU pool force production (i.e. models of dynamic force control through muscle excitation rather than static curve fitting): a pure feedforward and a combined feedforward-feedback configuration. For both strategies, our aim is to achieve online control for novel tasks (i.e. tasks for which the controller has not been optimised). While the present work is technically motivated, answering these questions can also shed light on the functional role of the large range of MU pool structures observed in human muscles (see [10]).

## 2 Models

For the present investigation, muscle force production is assumed to take place under feedforward control with an optional feedback element (Figure 1). Tasks are specified using a desired force signal and a FVL trajectory, which is a single gain signal that combines the muscle’s length and contraction speed state and the effect of state on the muscle’s force generating capacity. The task is communicated to *N* parallel MUs through a common neural drive, which drives recruitment and rate coding, resulting an excitation impulse train for each MU. The excitation impulses trigger the activation dynamics of each MU, leading to a total muscle force as a sum of the contraction forces of the *N* MUs. In the present work, we compare nine different MU pool models, which differ in terms of their recruitment and rate coding strategies, whereas the muscle fibres in each pool convert excitation to force in an identical manner, as detailed below. Additionally, we compare two control strategies for neural drive computation: pure feedforward and a combination of feedforward and feedback.

**Figure 1:**
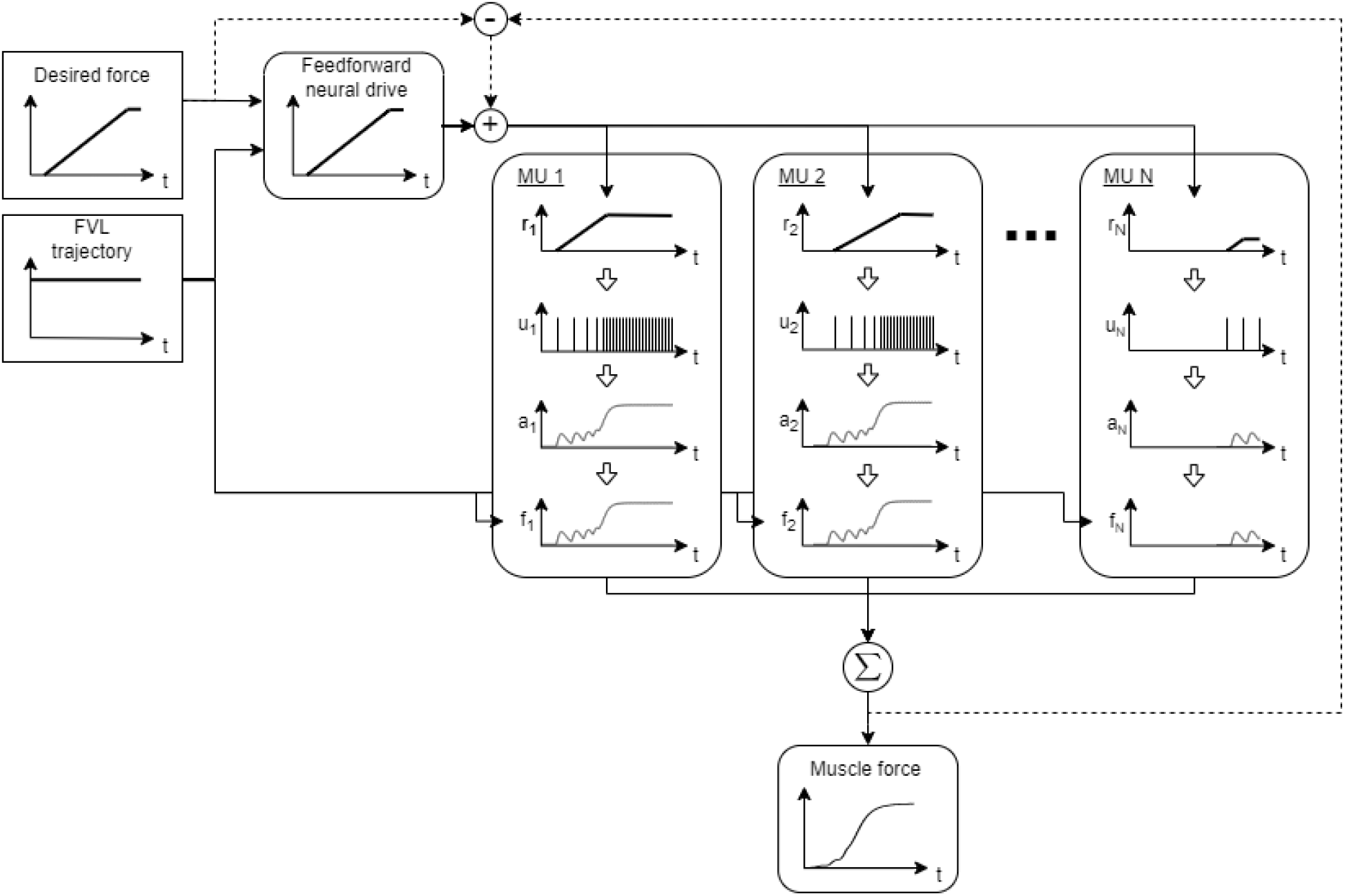
A schematic illustration of force generation in the multi-motor-unit muscle models. Time- based desired force and force-velocity-length (FVL) signals together with the optional feedback error signal (dashed line) are used to compute neural drive, which governs recruitment, rate coding, and force generation in *N* parallel motor units (MU). The *i*^th^ MU (*i* = 1, …, *N*) converts the neural drive to firing rate *r*_*i*_, which is used to compute the excitation impulse train *u*_*i*_. The excitation *u*_*i*_ triggers changes in the activation state *a*_*i*_, which, together with the MU’s isometric strength and FVL state, determine its force output *f*_*i*_.

### 2.1 Neural drive computation

The neural drive signal *I*(*t*) at time *t* in Figure 1 comprises feedforward and feedback elements, *I*_*ff*_ (*t*) and *I*_*fb*_(*t*), respectively,

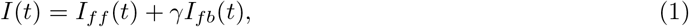

where setting the feedback gain *γ* to zero disables feedback. Although the two components are not intended to replicate specific neural circuits, *I*_*ff*_ (*t*) can be taken to generally represent a motor plan generated by the central nervous system, whereas *I*_*fb*_(*t*) represents reflex-like modifications to the plan during task execution. Since *I*(*t*) does not directly represent a physical variable, its units can be chosen freely. For the ease of interpreting neural drive as an intermediate force control signal, *I*(*t*) and its components are considered force-like with units of Newtons.

For an ideal motor plan, the mapping from desired force *F*_*d*_(*t*) at time *t* to *I*_*ff*_ (*t*) should be the inverse of the mapping from *I*(*t*) to the total muscle force *F*_*M*_(*t*). This would produce perfect match between *F*_*d*_(*t*) and *F*_*M*_(*t*) with feedback only needed in case of external perturbations. However, given the complex activation and FVL dynamics of MU pools, *F*_*M*_(*I*(*t*)) can only be obtained through simulations. This rules out direct curve-fitting of *F*_*M*_(*t*) to *F*_*d*_(*t*), and it also makes obtaining the inverse *I*(*F*_*M*_(*t*)) for a functionally relevant range of operation unfeasible, particularly for non-steady-state tasks. Instead, a simple mapping for *I*_*ff*_ (*t*) is constructed using a constant baseline muscle tone and *F*_*d*_(*t*) normalised with the FVL gain *g*(*t*) and with a delay *τ* between the input and output. The resulting mapping

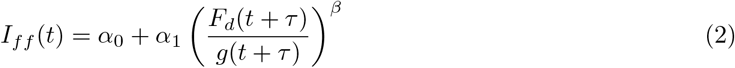

has three parameters, *α*_0_, *α*_1_, and *β*, in addition to the delay. The feedback component of the neural drive is given by

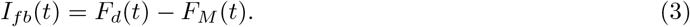

Given the role of the neural mapping in inverting the MU pool dynamics, and our goal of achieving control without reliance on task-specific optimisation, the mapping parameters are assumed to depend on the MU pool but to be common for all tasks the MU pool performs. The parameter values are obtained by optimisation in two stages. First, the feedforward parameters *α*_0_, *α*_1_, *β* and *τ* are optimised with feedback disabled (i.e. *γ* = 0). Second, the feedforward parameters are kept constant and *γ* is optimised. A set of elementary tasks (see Section 2.4.1) is used for the optimisations and the objective is to minimise the sum of average square errors

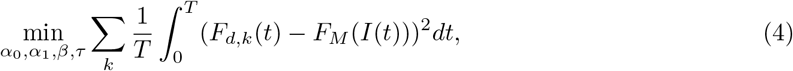

where *k* is the index for the elementary tasks.

### 2.2 Motor unit pool models

#### 2.2.1 Recruitment and rate coding strategy

Recruitment and rate coding strategy governs how the firing rate *r*_*i*_ of MU *i* changes with the desired force *F*_*d*_(*t*). In phenomenological MU pool models, it comprises two components: i) a threshold function, which determines the recruitment thresholds *δ*_*i*_ such that *r*_*i*_ *>* 0 only when *I*(*t*) ≥ *δ*_*i*_, and ii) a rate function, which describes how *r*_*i*_ changes as *I*(*t*) increases beyond *δ*_*i*_. There are multiple ways to construct and parameterise such models (e.g. [13, 17, 23]), but we focus on three types of MU pool models which differ in their recruitment threshold functions and the general form of their rate functions. For each type of MU pool, we consider three variations in handling the maximum firing rate.

The three types of MU pool models are exponential-linear, matched-linear and mixed-logarithmic:

##### 1. Exponential-linear

This MU pool type utilises exponentially assigned recruitment thresholds inspired by [13] with a constant recruitment bandwidth Δ_*i*_, which is the range of neural drive values over which the MU force increases from minimum to maximum,

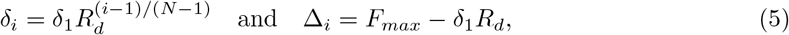

where *F*_*max*_ is the maximum isometric strength of the muscle. The shape of the threshold function is defined by the threshold for the first MU, *δ*_1_, and the (dimensionless) range of recruitment thresholds *R*_*d*_ = *δ*_*N*_ */δ*_1_ *>* 0.

Once a MU has been recruited (i.e. *I > δ*_*i*_), its firing rate *r*_*i*_ is determined by a capped linear rate function (e.g. [13, 17])

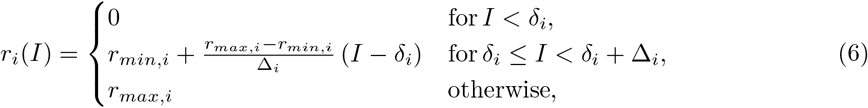

where *r*_*i*_ increases linearly from minimum firing rate *r*_*min,i*_ to maximum firing rate *r*_*max,i*_ over the recruitment bandwidth. Although this rate function does not capture some known effects, such as the tendency for a steeper initial slope [17, 23], it represents the gross firing rate behaviour with minimal parameters.

##### 2. Matched-linear

In this MU pool type, the recruitment bandwidth of MU *i* is matched to its isometric strength *f*_*max,i*_ with constant overlap between adjacent recruitment bands, leading to

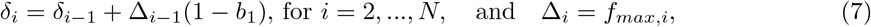

for a selected *δ*_1_. The dimensionless overlap parameter *b*_1_ ∈ [0, 1] represents the proportion of Δ_*i*−1_ where the forces produced by MUs *i* − 1 and *i* are simultaneously increased through rate coding. For a given desired range of recruitment thresholds *b*_1_ = 1 − (*δ*_*N*_ − *δ*_1_)*/*(*F*_*max*_ − *f*_*max,N*_) .

The linear, capped rate function (Eq. 6) is used for this MU pool type.

This MU pool type is included as a “linearised” version of the exponential-linear type, rather than as a realistic representation of human MU pools. That is, the steady-state tetanic force output from matched-linear pools increases roughly linearly with neural drive, which is not true for the other pool types, and hence may lead to different controllability properties.

##### 3. Mixed-logarithmic

This MU pool type uses a mixture of linear and exponential recruitment thresholds inspired by [12]

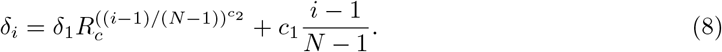

The shape of the threshold function is defined by the dimensionless parameters *R*_*c*_ and *c*_2_, and the force-like parameter *c*_1_ (units: Newtons). A given range of thresholds can be achieved by setting *R*_*c*_ = (*δ*_*N*_ − *c*_1_)*/δ*_1_.

The rate function for this MU pool type is the steady-state relationship arising from a single-compartment leaky-integrate-and-fire model of the motor neuron membrane [12], adjusted to utilise neural drive *I*(*t*),

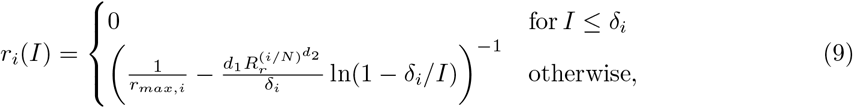

where the parameters *d*_2_ and *R*_*r*_ are dimensionless while *d*_1_ has the units of Ns, and ln() is the natural logarithm.

Multiple different ways to characterise the maximum firing rate *r*_*max,i*_ have been proposed (e.g. [13, 17, 23]) to account for the variability in experimental data [10] while also explaining phenomena such as onion-skin recruitment. To investigate the effects of variations in maximum firing rates within the MU pools, the three MU pool types are each combined three maximum firing rate schemes: i) constant *r*_*max*_ for all MUs

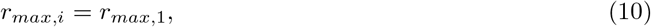

ii) linearly decreasing *r*_*max*_ from smallest to largest MU

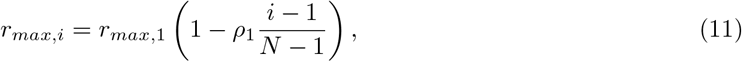

and iii) exponentially decreasing *r*_*max*_

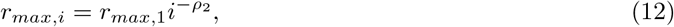

each defined for *i* = 2, …, *N* using the maximum firing rate for the first MU, *r*_*max*,1_ and the parameters *ρ*_1_ and *ρ*_2_, for the linear and exponential schemes, respectively. The resulting nine different MU pool models are summarised in Table 1.

**Table 1:** Motor unit pool models: combinations of threshold and rate functions and maximum firing rate schemes, and the abbreviations used in figures.

| Threshold function | Rate function | $r_{max}$ scheme | Model abbr. |
| --- | --- | --- | --- |
| Exponential, Eq. 5 | Linear, Eq. 6 | Constant, Eq. 10 | exp-lin const |
|  |  | Linear, Eq. 11 | exp-lin lin |
|  |  | Exponential, Eq. 12 | exp-lin exp |
| Strength matched, Eq. 7 | Linear, Eq. 6 | Constant, Eq. 10 | matched-lin const |
|  |  | Linear Eq. 11 | matched-lin lin |
|  |  | Exponential, Eq. 12 | matched-lin exp |
| Mixed, Eq. 8 | Logarithmic. Eq. 9 | Constant, Eq. 10 | mixed-log const |
|  |  | Linear, Eq. 11 | mixed-log lin |
|  |  | Exponential, Eq. 12 | mixed-log exp |

#### 2.2.2 Excitation impulse train construction

Once task *F*_*d*_(*t*) and *g*(*t*) have been converted to *I*(*t*) and then to *r*_*i*_(*t*) for all MUs in the pool, excitation impulse trains for each MU can be constructed. This is done by identifying the first instants when the time since last impulse exceeds the desired inter-impulse period 1*/r*_*i*_(*t*). The first impulse for MU *i* is fired at *t* = *ϕ*_*i*,1_ which corresponds to the first instance in the simulation when *r*_*i*_(*t*) *>* 0. The times *ϕ*_*i,j*_ for the following impulses *j* = 2, … are determined one by one by

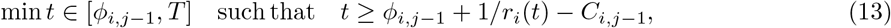

where *T* is the duration of the simulation and *C*_*i,j*_ is a corrective term accounting for the error introduced by discrete time, i.e. the fact that the equality in Eq (13) can only hold for a limited set of *r*_*i*_ values during simulations. This correction is calculated using linear interpolation to estimate when the *j*^th^ impulse would have been fired in continuous time, resulting in

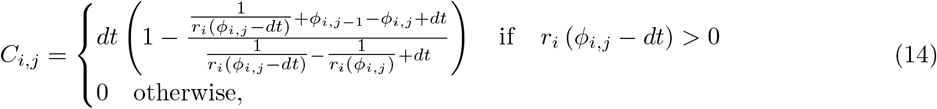

where *dt* is the simulation time step. The excitation impulse train is then

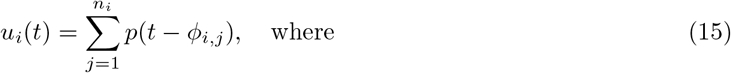

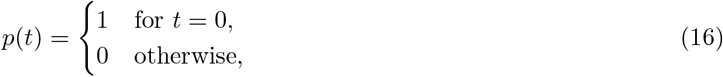

and *n*_*i*_ is the number of impulses for MU *i* during the simulation.

#### 2.2.3 Motor unit force production

The total muscle force *F*_*M*_(*t*) produced by a pool of *N* MUs is

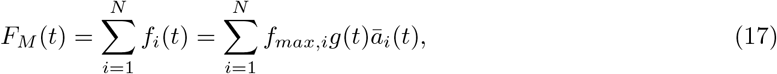

where the force *f*_*i*_(*t*) produced by the *i*^th^ MU depends on its maximum isometric strength *f*_*max,i*_, FVL state *g*(*t*) as well as its normalised activation state *ā* _*i*_(*t*).

Within the MU pool, the index *i* = 1, ‥, *N* orders the MUs by size from smallest to largest, with isometric strengths, following [13],

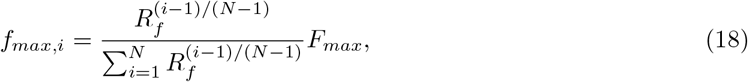

where *R*_*f*_ = *f*_*max,N*_ */f*_*max*,1_ is the range of MU strengths and *F*_*max*_ is the isometric strength of the entire muscle. Note that the indexing (*i*− 1)*/*(*N* − 1) is used to define MU characteristics for ease of parameter interpretation. Given that *N* is large, the difference compared to the indexing *i/N* used in the original MU pool [13] is negligible.

For the purposes of the present work, the FVL properties are condensed into a time-dependent trajectory *g*(*t*), which is defined externally in the task specifications (see Section 2.4.1) and identical for all MUs. Note, however, that *g*(*t*) in the tasks arises from changes in a muscle’s length and contraction speed. Hence, we are inherently assuming that all MUs have identical normalised length and contraction speed trajectories, and that their FVL functions are also the same and, for simulated task data, match the Hill-type model properties underlying the FVL trajectory data.

The activation dynamics (i.e. excitation-contraction coupling) for each MU are described by *ā* _*i*_(*t*) = *a*_*i*_(*u*_*i*_(*t*), *t*)*/a*_*max,i*_ ∈ [0, 1]. The unnormalised activation state *a*_*i*_(*u*_*i*_(*t*), *t*)) follows third order activation dynamics [39], which captures both realistic twitch responses and their nonlinear summation, and it is driven by an excitation impulse train *u*_*i*_(*t*). The normalisation factor *a*_*max,i*_ corresponds to the unnormalised tetanic activation level when the excitation impulses are fired at maximum rate *r*_*max,i*_. For the present work, we assume that the activation dynamics of all MUs are identical, that is, that twitch properties do not vary between MUs except for the twitch amplitude, which depends on *f*_*max,i*_.

Note that the assumptions made about identical contractile properties for all MUs are simplifications. Experimental data suggests that strains and strain rates are not homogeneous across muscles [40] and different muscle fibre types can differ in properties such as maximum contraction speed [41] and the shape of the twitch pulse [39]. However, to retain our focus on the control aspects of MU pool structures (i.e. recruitment and rate coding) and to limit the number of confounding factors, we consider variations in muscle fibre properties outside the present scope.

### 2.3 Motor unit pool parameters

The common parameters of the MU pools (*N, F*_*max*_, *r*_*max*,1_, *R*_*f*_, *δ*_1_, *δ*_*N*_, and activation dynamics characterised by time-to-peak and half-relaxation time) are adjusted to represent the two muscles studied, TA and SF (Table 2). For the TA, the MU pools are parameterised to represent the entire muscle with *N* = 400 MUs based on [12], *F*_*max*_ = 400 N picked from middle of the range reported by [42] and MU strength range set to *R*_*f*_ = 100 [13]. For the SF, the MU pools only represent the *N* = 200 smallest MUs whose total strength adds up to *F*_*max*_ = 200 N, rather than a larger pool representing the entire muscle. Only this subsection of the pool was needed as the peak forces in the SF tasks (see Section 2.4.1) are well below the recruitment threshold of the muscle’s larger MUs. The *R*_*f*_ and *δ*_*N*_ values for the SF are set to reflect this partial pool architecture. The twitch parameters for the TA were set to fall within the ranges reported by [43, 44], and for the SF, they were set to match the reaching simulation data [38].

**Table 2:** MU pool characteristics for tibialis anterior and a generic shoulder flexor. The sources of values are discussed in Section 2.3.

| Parameter | Tibialis anterior (TA) | Shoulder flexor (SF) |
| --- | --- | --- |
| $N$ | 400 | 200 |
| $F_{max}$ (N) | 400 | 200 |
| $R_f$ | 100 | 10 |
| Time-to-peak (ms) | 60 | 70 |
| Half-relaxation time (ms) | 70 | 80 |
| $\delta_1$ (N) | 0.5 | 0.5 |
| $\delta_N$ (N) | $0.5F_{max}$ | $0.95F_{max}$ |
| $r_{max,1}$ (Hz) | 25 | 50 |

To facilitate easy comparison, the range of recruitment thresholds, represented by *δ*_1_ and *δ*_*N*_, is kept constant for all MU pools representing the same muscle (see Figure 2 for TA). Based on the muscle specific parameters, the threshold function parameters are adjusted accordingly: *R*_*d*_ = 400 for TA and *R*_*d*_ = 380 for SF, *b*_1_ ≈ 0.496 for TA and *b*_1_ ≈ 0.041 for SF, *c*_1_ = 29.195 N and *c*_2_ = 1.833 for both muscles based on [12], and *R*_*c*_ ≈ 341.6 for TA and *R*_*c*_ ≈ 321.6 for SF.

**Figure 2:**
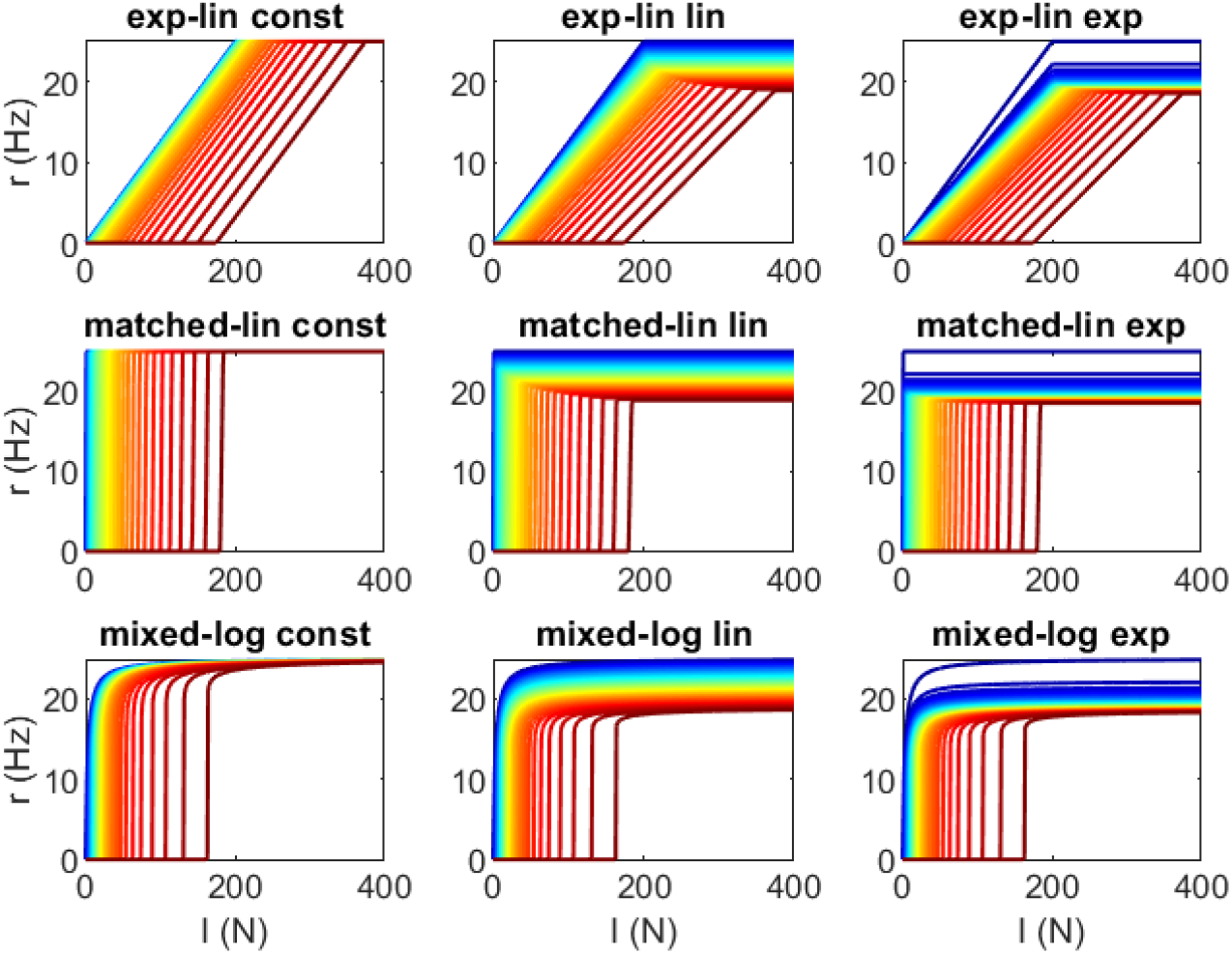
Firing rate functions for the nine MU pool models of TA. Firing rates (*r*) vs. neural drive (*I*) for the MUs in pools representing the TA for the different recruitment schemes. The line color varies from dark blue for the first MU to dark red for the last MU. Only every 10^th^ of the 400 MUs are shown.

The minimum firing rate *r*_*min,i*_ in the linear rate function is assumed to be zero for easy comparison with the logarithmic rate function, for which *r*_*min*_ = 0 implicitly. While this assumption does not match known MU behaviour at the recruitment threshold (e.g. [17]), the difference is only notable if the neural drive is maintained at a MU’s recruitment threshold for long enough to evoke multiple excitation spikes, which is only true for a few MUs in a pool even during long, sustained contractions.

The maximum firing rate *r*_*max*,1_ is set to 25 Hz for TA based on [5, 12] to ensure comparability with reference data and to 50 Hz for SF to account for existing data on deltoid during slowly varying contractions [45] and the general observation that rapid contractions can elicit significantly higher firing rates than slower ones [46]. The parameters for the logarithmic rate function are adapted from [12] to *d*_1_ =0.1 Ns, *d*_2_ =1.47, and *R*_*r*_=2.4. The *r*_*max*_ scheme parameters are set to *ρ*_2_ = 0.05 based on [12] and to *ρ*_1_=0.25 to achieve roughly equal reduction in *r*_*max*_ in the linear and exponential schemes.

### 2.4 Simulations

#### 2.4.1 Tasks

Each task for the simulations is defined as a pair of desired force *F*_*d*_(*t*) and FVL trajectory *g*(*t*) for time *t* ∈ [0, *T*]. Optimisation of the parameters of the neural drive (Eq. 2) is done using a set of scaled, artificially constructed elementary tasks. The performance of the MU pool models is then evaluated using tasks based on either experimental or musculoskeletal simulation data.

The task sets for optimisation are constructed using three elementary tasks (Figure 3A): an isometric contraction, and concentric and eccentric contractions where the muscle length follows the minimum-jerk trajectory [47]. In all of these tasks, the target force profile is parabolic, and it occurs during the first half of the movement for the concentric and isometric tasks and during the second half for the eccentric task. The amplitude of the target force is scaled to match the range of forces required during experimental or MSK simulation test tasks. This range is achieved using two levels of force amplitude for each muscle, 0.3*F*_*max*_ and 0.5*F*_*max*_ for the TA and 0.05*F*_*max*_ and 0.2*F*_*max*_ for the SF (Nb. Figure 3A shows the normalised force target before scaling), resulting in a total of six tasks used to optimise the neural drive parameters for each muscle. The duration of all of the tasks is 2 sec, with target force production occurring over 1 sec. All contractions start at optimal muscle length and the change in muscle length is *±*20% for the concentric/eccentric tasks. FVL gains are computed from length and contraction speed using the Hill-type muscle model used in simulations of the test tasks for the SF [38].

**Figure 3:**
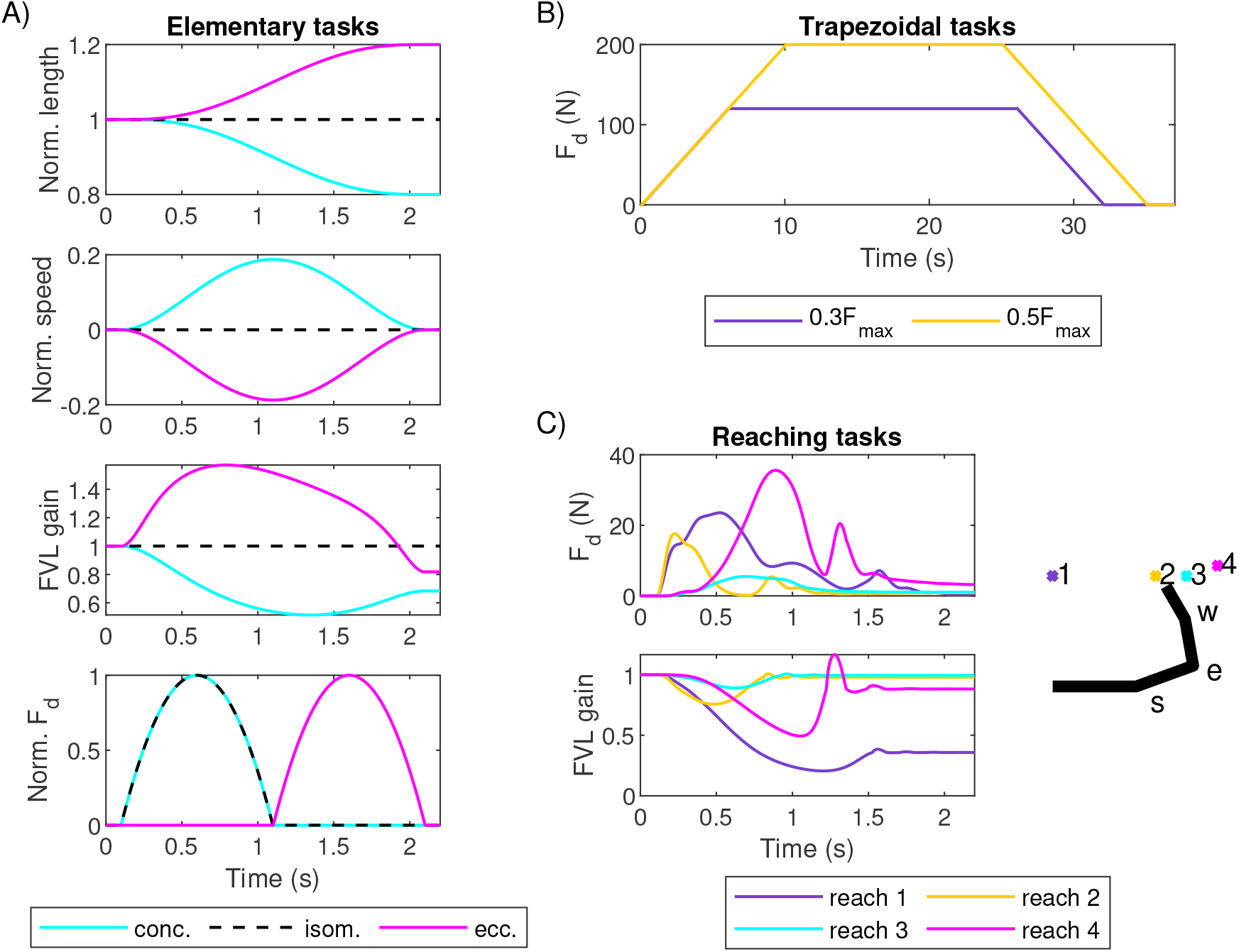
Force matching tasks. (A) Elementary concentric, isometric and eccentric tasks used to optimise neural drive. From the top, panels show normalised muscle length, normalised contraction speed, the resulting force-velocity-length (FVL) gain, and normalised desired force. (B) Trapezoidal tasks with two different force amplitudes. Only desired force is shown as the tasks are isometric. (C) Four reaching tasks with desired forces and FVL gains. Target locations for the reaches are indicated by a top-down view of the workspace and the initial position of (right) arm where s, e, and w indicate shoulder, elbow and wrist joints, respectively.

Once the parameters for the neural drive mapping for each MU pool have been obtained, the ability of the MU pools to match functional force profiles is tested. The test tasks for the MU pools representing the TA are slow isometric trapezoidal force profiles with plateaus at 30% and 50% *F*_*max*_ matching the experimental conditions of data from [5] (Figure 3B). For the SF, the test tasks are four simulated force profiles with matched FVL trajectories produced by a MSK model of the human arm [38], representing a group of reaching movements where the SF functions as an agonist (Figure 3C).

#### 2.4.2 Matching errors

Three errors are used to quantify the ability of the MU pool models to match desired force profiles. Each error emphasises a different aspect of the match, and they reflect the functionally different contexts in which force production could occur. First, the root-mean-square error

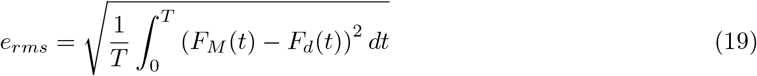

reflects the average match between target and muscle force during each task, and it serves as an overall measure of success. Second, the maximum absolute error normalised with the peak force amplitude for the task

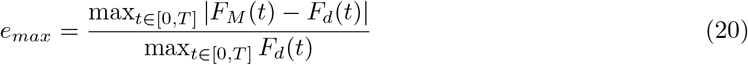

measures the worst instantaneous match. Finally, the total error

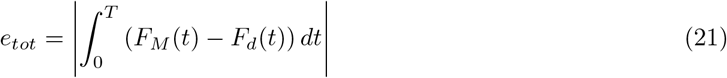

captures how much the impulse delivered over the entire task differs from the target. In a freely moving constant-load system, *e*_*tot*_ is proportional to the velocity error at the end of the task, and, unlike *e*_*rms*_, it does not penalise a muscle for compensation, that is, for force below the target during one part of the task but exceeding the target during another part.

## 3 Results

### 3.1 Neural drive mapping

Optimisation of neural drive parameters for the elementary tasks resulted in mappings from desired force *F*_*d*_ to feedforward neural drive *I*_*ff*_ which differ mainly between muscles and MU pool types, with little differences shown between different *r*_*max*_ schemes (Figure 4, left panels). While the mappings for matched-linear type pools are approximately linear, the other pools exhibit non-linearity, particularly when *F*_*d*_ (or *F*_*d*_*/g* for non-isometric contractions) is low. The neural drive mappings largely cancel out the nonlinearity inherent in the tetanic response of the MU pools at different levels of *I* (Figure 4, middle panels), resulting in a nearly linear combined mappings from *F*_*d*_ to tetanic *F*_*M*_(Figure 4, right panels). Note that Figure 4 represents isometric, steady-state contractions and does not capture the dynamic changes in *F*_*M*_arising from changes in *F*_*d*_ or the muscle’s FVL state.

**Figure 4:**
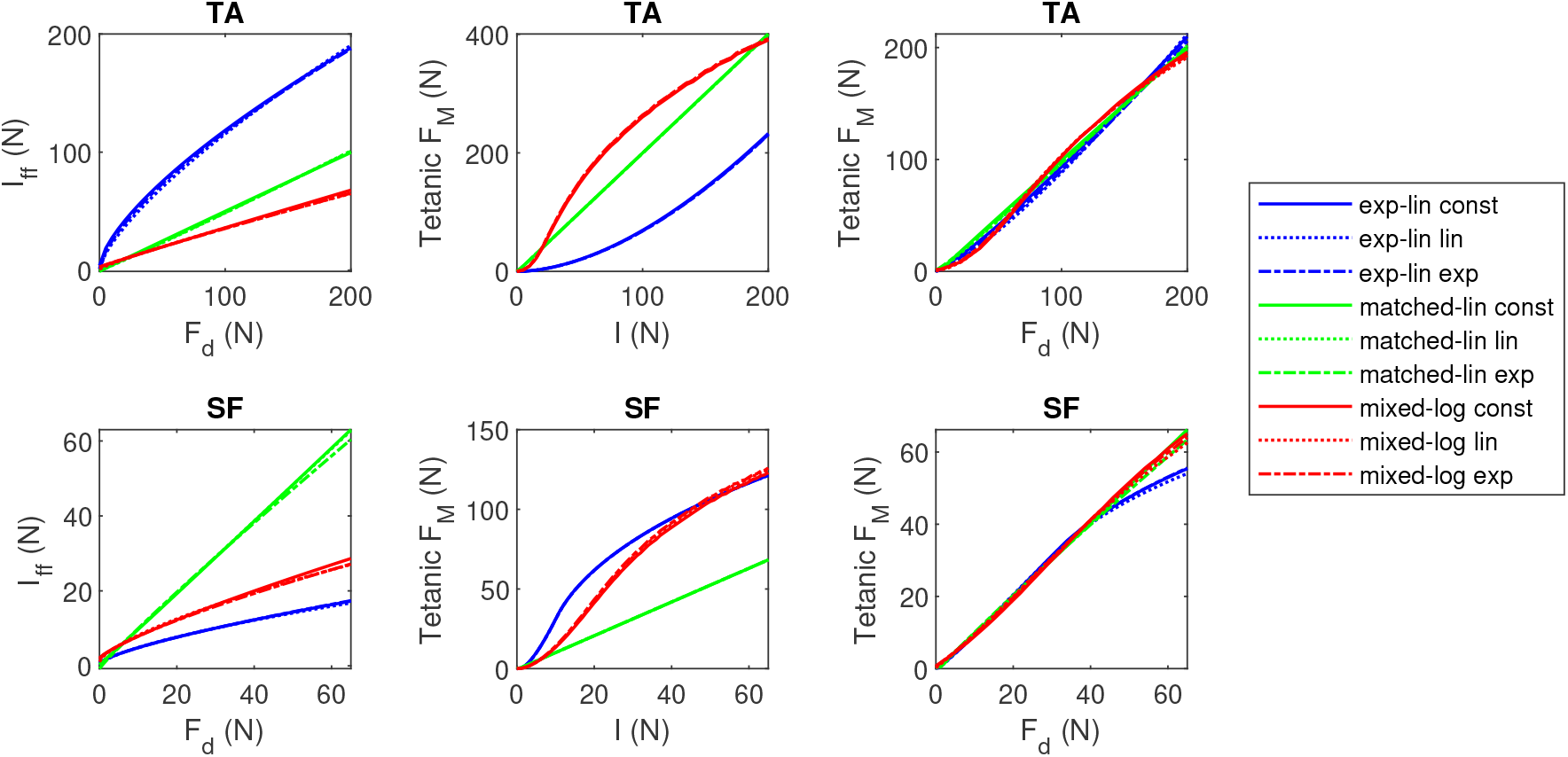
The mappings from desired force (*F*_*d*_) to feedforward neural drive (*I*_*ff*_) to isometric tetanic muscle force (*F*_*M*_) for the nine MU pool models representing TA and SF. The top row represents TA and bottom row SF, with panels on reach row showing from left to right: mapping from *F*_*d*_ to *I*_*ff*_ after neural drive parameters have been optimised for each MU pool and muscle, mapping from *I* = *I*_*ff*_ (i.e. feedforward only) to tetanic *F*_*M*_, and a combined mapping from *F*_*d*_ to tetanic *F*_*M*_. Isometric contraction at optimal length is assumed (*g* = 1).

### 3.2 TA/trapezoidal tasks under feedforward control

All MU pools parameterised for the TA matched the trapezoidal target forces generally well under feedforward control (Figure 5). However, there are distinct differences in the accuracy of the plateau match and minor differences during the ascending and descending force ramps. The matched-linear type pools match the desired force profile the best when performance is measured using *e*_*rms*_ and *e*_*max*_ (Figure 6A), with the matched-linear pool with constant *r*_*max*_ producing the best matching overall. The plateau force for all of the matched-linear pools was, on average, within 2.7 N of the target on both trapezoidal tasks. The exponential-linear pools and the mixed-logarithmic pools produced more varied results, with different error measures favouring different pools at different tasks. In terms of plateau force match, the exponential-linear pools produced sustained contraction, on average, 4.2-8.6 N below the lower 120 N target and 5.9-12.2 N above the higher 200 N target. The effect was opposite for the mixed-logarithmic pools, which produce 4.5-5.2 N in excess of the lower target and 4.0-8.8 N below the higher target. The exponential-linear pools also display larger force variability during the plateau compared to the other models, although this variability is comparable to human performance (see [5]).

**Figure 5:**
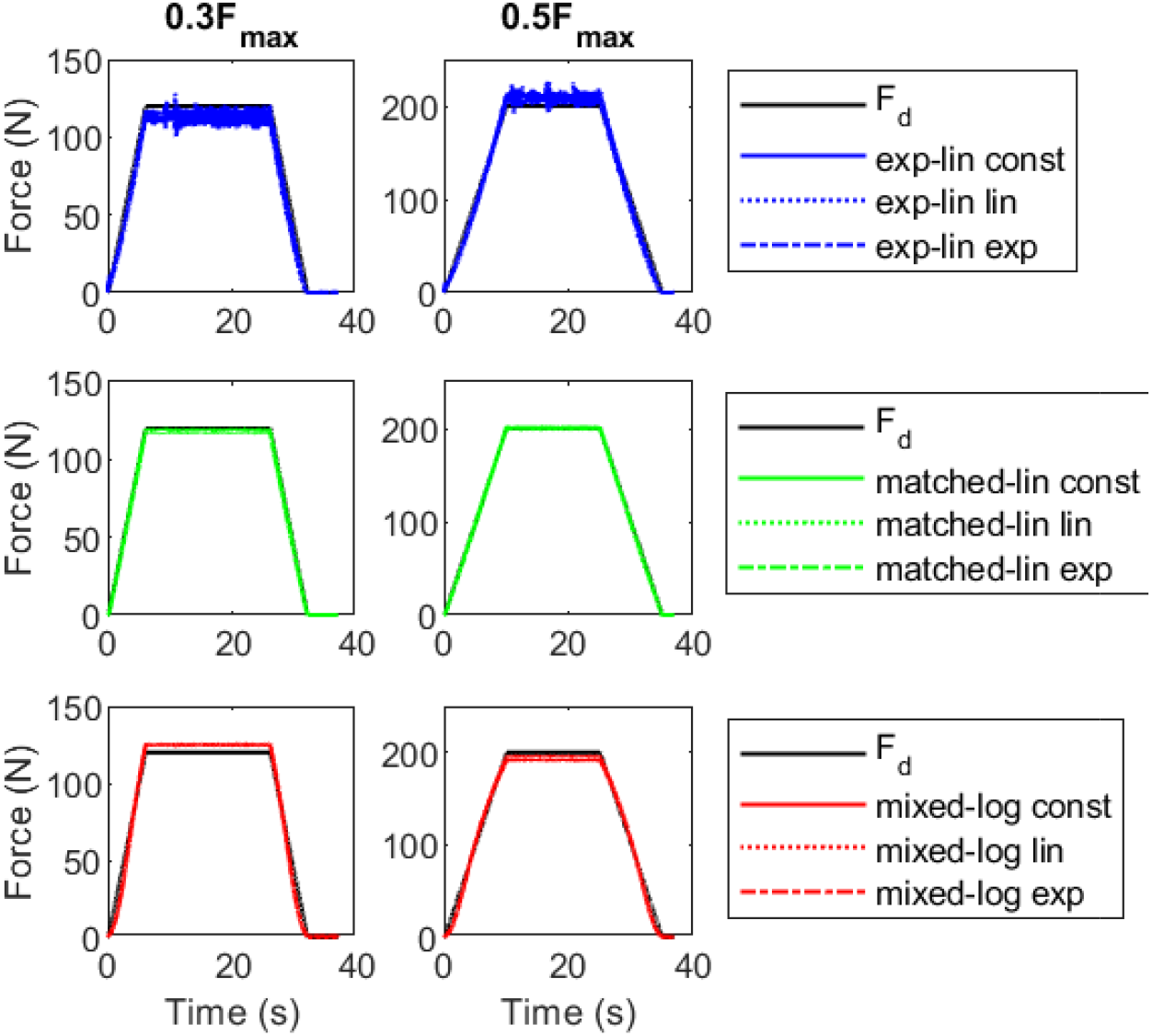
Force match for the trapezoidal tasks by TA for pure feedforward control. Panel columns show the trapezoidal tasks at the two force levels. The force curves have been grouped by basic MU pool type.

**Figure 6:**
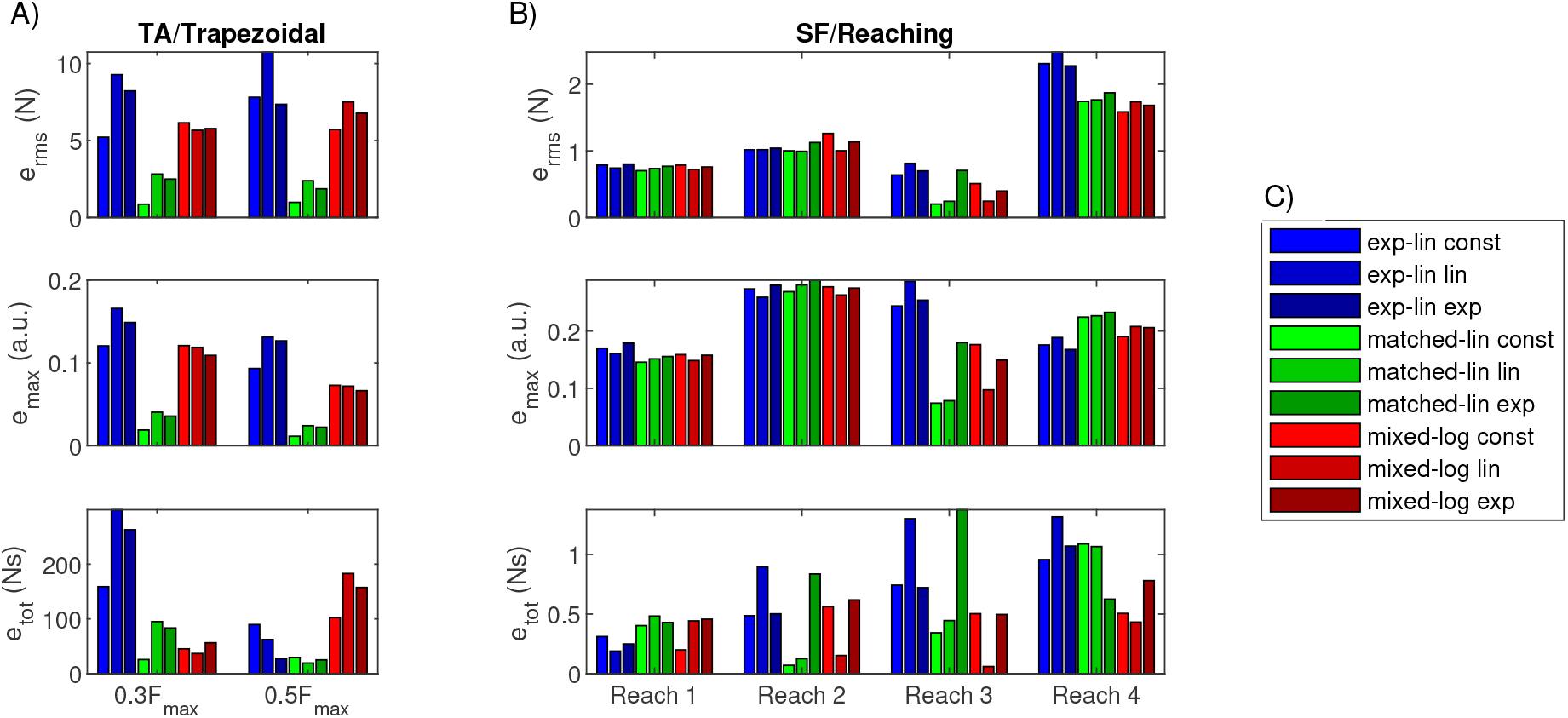
Force match errors in trapezoidal and reaching tasks for pure feedforward control. (A) Performance of the nine MU pool models of the TA in two trapezoidal tasks. (B) Performance of the nine MU pool models of the SF in four reaching tasks. (C) Legend for the MU pool colours in panels A and B.

No *r*_*max*_ scheme appeared to universally improve the ability of all MU pools to match the trapezoidal forces. However, constant *r*_*max*_ leads to favourable outcomes for both exponential-linear and matched- linear pool types, particularly in the low force task. More prominently, the *r*_*max*_ scheme affects the firing rate patterns observed in the MUs during the task, as shown in Figure 7A for the 0.3*F*_*max*_ task (see Supplementary Figure 1 for the higher force task), which displays the true instantaneous firing frequencies extracted from the excitation signals *u*_*i*_ and smoothed using a 5-sample moving average filter. There are two main differences in the firing rate behaviors of the MU pools. First, firing rates in the exponential-linear pools increase and decrease slowly during force ramping, whereas MUs in the other pools achieve their steady-state firing rates nearly instantaneously. Qualitative comparison with firing rates extracted from high-density surface EMG data [5] during the same task (Figure 7B for three examples, see Supplementary Figure 2 for full dataset) suggests that the slower rate increase matches real MU behaviour better, although some MUs, particularly those recruited late in the task, develop their maximum firing rates very fast. Second, the range of maximum firing rates developed during the task vary by both MU pool type and by *r*_*max*_ scheme. While the variation in the maximum firing rate in the matched-linear pools can be attributed nearly solely to the *r*_*max*_ scheme, in the other pool types, MUs recruited later in the task can exhibit lower firing rates both due to decreasing *r*_*max,i*_ and due to submaximal recruitment (i.e. *r*_*i*_ *< r*_*max,i*_). The experimental data (Supplementary Figure 2) suggest that differences in the maximum firing rate of MUs and the pattern of lower firing rates in later recruited MUs are common but not always present in real muscle contractions.

**Figure 7:**
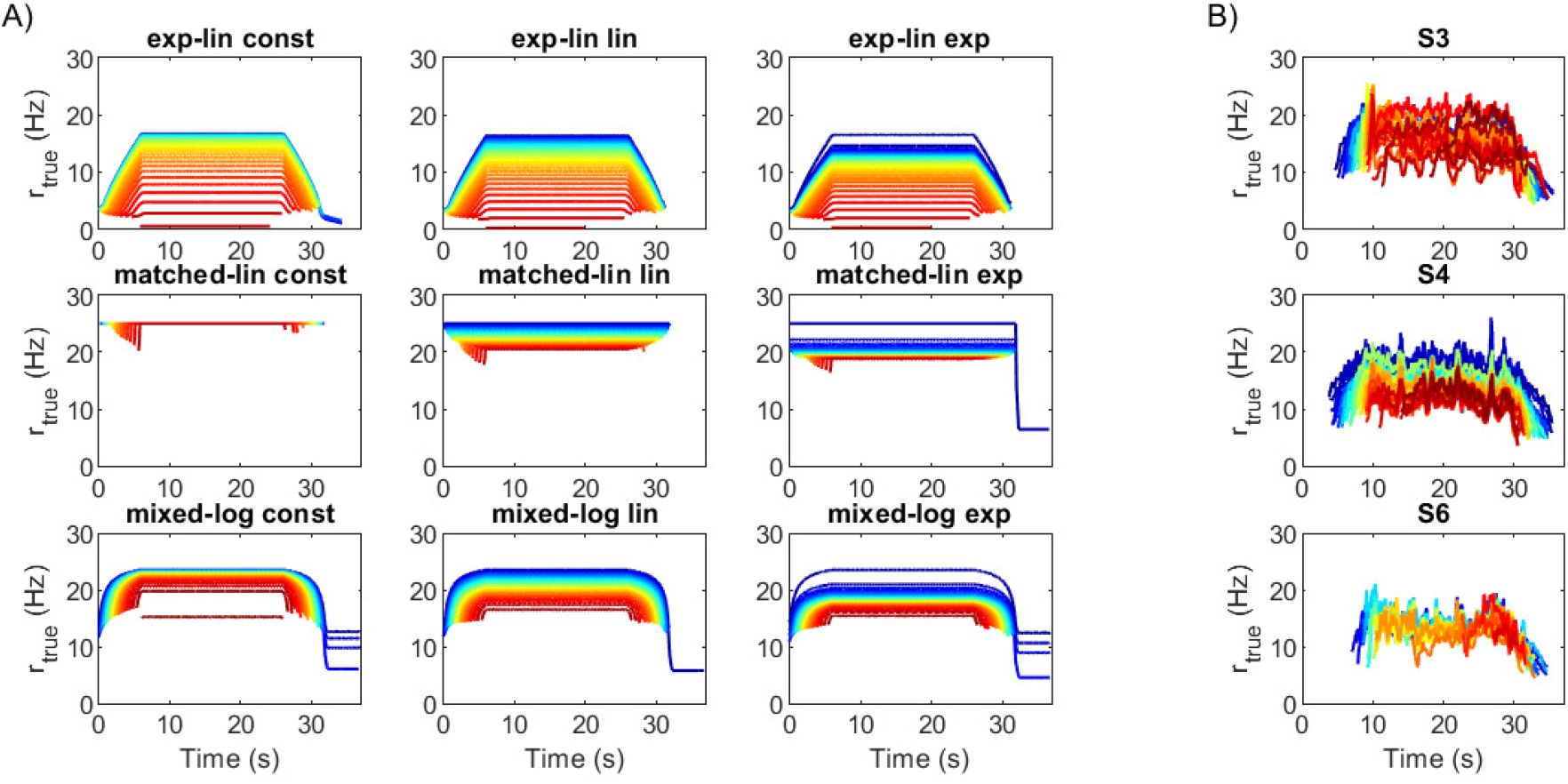
True firing frequencies for the TA/trapezoidal 0.3*F*_*max*_ task for pure feedforward control. (A) Simulated frequencies extracted from excitation impulse trains for the nine MU pools. (B) Examples (subjects S3, S4 and S6) of experimental data for the same task [5]. The colours in all panels indicate order of MU recruitment from first (dark blue) to last (dark red). Only every 10^th^ MU is shown in A. Note that there is no correspondence between the ordering of panels in A and B.

Qualitative inspection of the experimental firing rate data from [5] also suggested that for some, but not all, test subjects, the 0.5*F*_*max*_ task elicited higher firing rates in the early recruited MUs than the 0.3*F*_*max*_ task (see Supplementary Figure 2). This behaviour cannot be replicated by the matched- linear and mixed-logarithmic pools without significant adjustment to the pool parameters; due to small recruitment bandwidths in these pool models, the firing rate of the smallest MUs reaches *r*_*max,i*_ at low levels of desired force. In contrast, all of the exponential-linear pools show an increase in the firing frequencies of the smallest MUs as force demand is increased from 0.3*F*_*max*_ to 0.5*F*_*max*_. Hence, the nine pool models fall grossly within the large range of behaviours exhibited by real muscles. However, the qualitative comparison suggests that the exponential-linear models produce firing rate patterns closest and the matched-linear models furthest from real muscle recruitment behaviour.

### 3.3 SF/reaching tasks under feedforward control

The performance of the MU pool models in representing a SF in the four reaching force matching tasks is shown in Figure 8 with corresponding error metrics displayed in Figure 6B. The overall error levels suggest that there is relatively little difference between the MU pool types, and as with the TA and trapezoidal tasks, the *r*_*max*_ scheme has relatively little impact on the overall performance, although performance varies by error and task. If the MU pools are ordered from lowest to highest error for each task, the matched-linear pool with constant *r*_*max*_ and mixed-logarithmic pool with linear *r*_*max*_ show a slight tendency to perform most consistently well.

**Figure 8:**
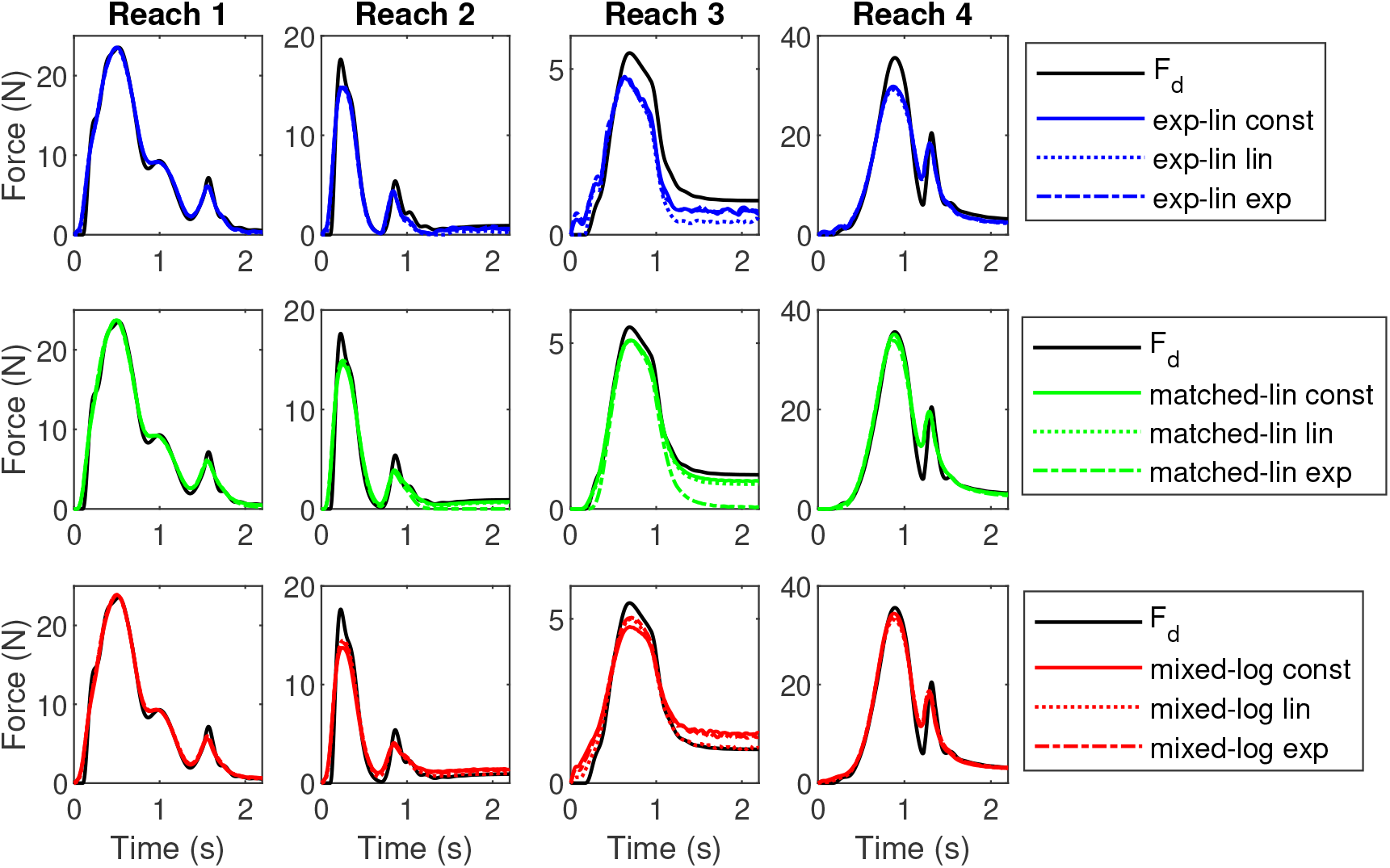
Force match for the simulated reaching tasks by SF for pure feedforward control. Panel columns show the four reaching tasks. The force curves have been grouped by basic MU pool type.

Inspection of the force signals (Figure 8) suggests that reaching tasks 3 and 4 elicited the most variable performance from the MU pools. The MU pool models struggle to match both the low peak force and the even lower sustained contraction of reach 3, likely at least partially due to the relatively small number of MUs recruited. While the exact number of MUs active during reach 3 varies by pool, for reference, the peak force corresponds roughly to the maximal contraction of the smallest 15 MUs in the pool and the sustained force to the smallest 3 MUs. In reach 4, the peak amplitude is less problematic, although not quite matched by the exponential-linear or, to a lesser extent, by the mixed-logarithmic pools. However, the rapid force decay followed by a second force peak in reach 4 proved difficult for the MU pools to match, with none of the pools reaching the desired local force minimum. Reaches 1 and 2 also have local force minima followed by secondary peaks, which the pools match largely successfully, so the problematic feature in reach 4 likely relates to the short time scale and timing of the force changes relative to the immediate history of MU firing.

### 3.4 Effect of feedback

The addition of feedback leads to improved performance in both the trapezoidal and reaching tasks for all pools using all error metrics with two exceptions (Figures 9–10). These exceptions were increases in all errors for both trapezoidal tasks for the matched-linear pool with constant *r*_*max*_, potentially suggesting failed optimisation of the feedback gain *γ*, and an increase in *e*_*tot*_ for reach 2 also for the same MU pool model. The addition of feedback benefited the physiologically motivated exponential-linear and mixed-logarithmic MU pool types more than the matched-linear pools. For the TA and trapezoidal tasks, this eliminated much of the differences in the MU pool models’ performance seen under pure feedforward control. For the SF and reaching tasks, the exponential-linear and mixed-logarithmic MU pools produce overall lower errors than the matched-linear pools when feedback is enabled, although task- and error-dependent variation in performance remains notable.

**Figure 9:**
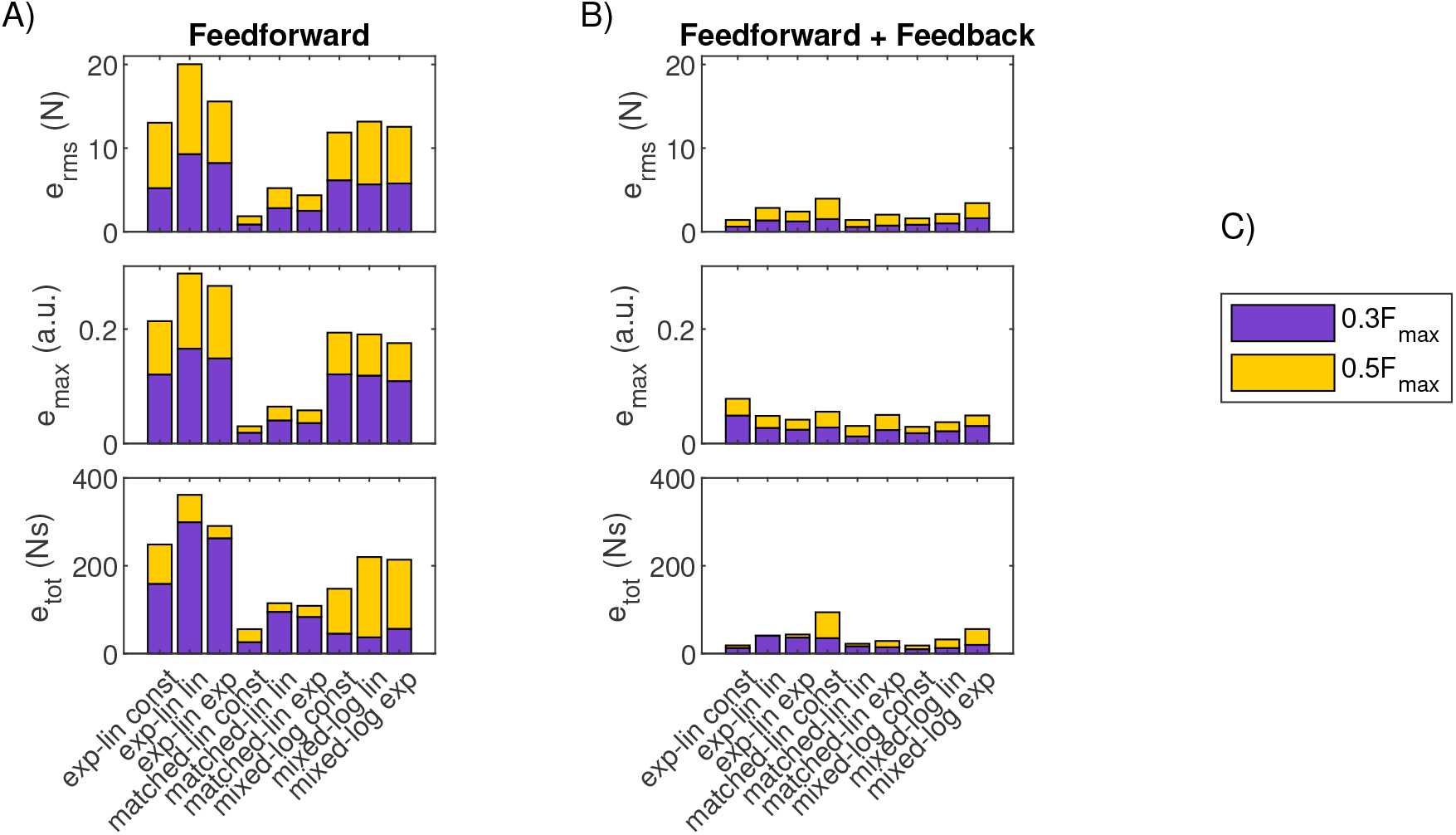
Force matching errors for the trapezoidal tasks for feedforward and combined feedforward-feedback control. (A) Stacked performance errors for the nine MU pool models of the TA for the two trapezoidal tasks under pure feedforward control. (B) Stacked performance errors for the nine MU pool models of the TA for the two trapezoidal tasks under combined feedforward-feedback control. (C) Legend for task colours in panels A and B.

**Figure 10:**
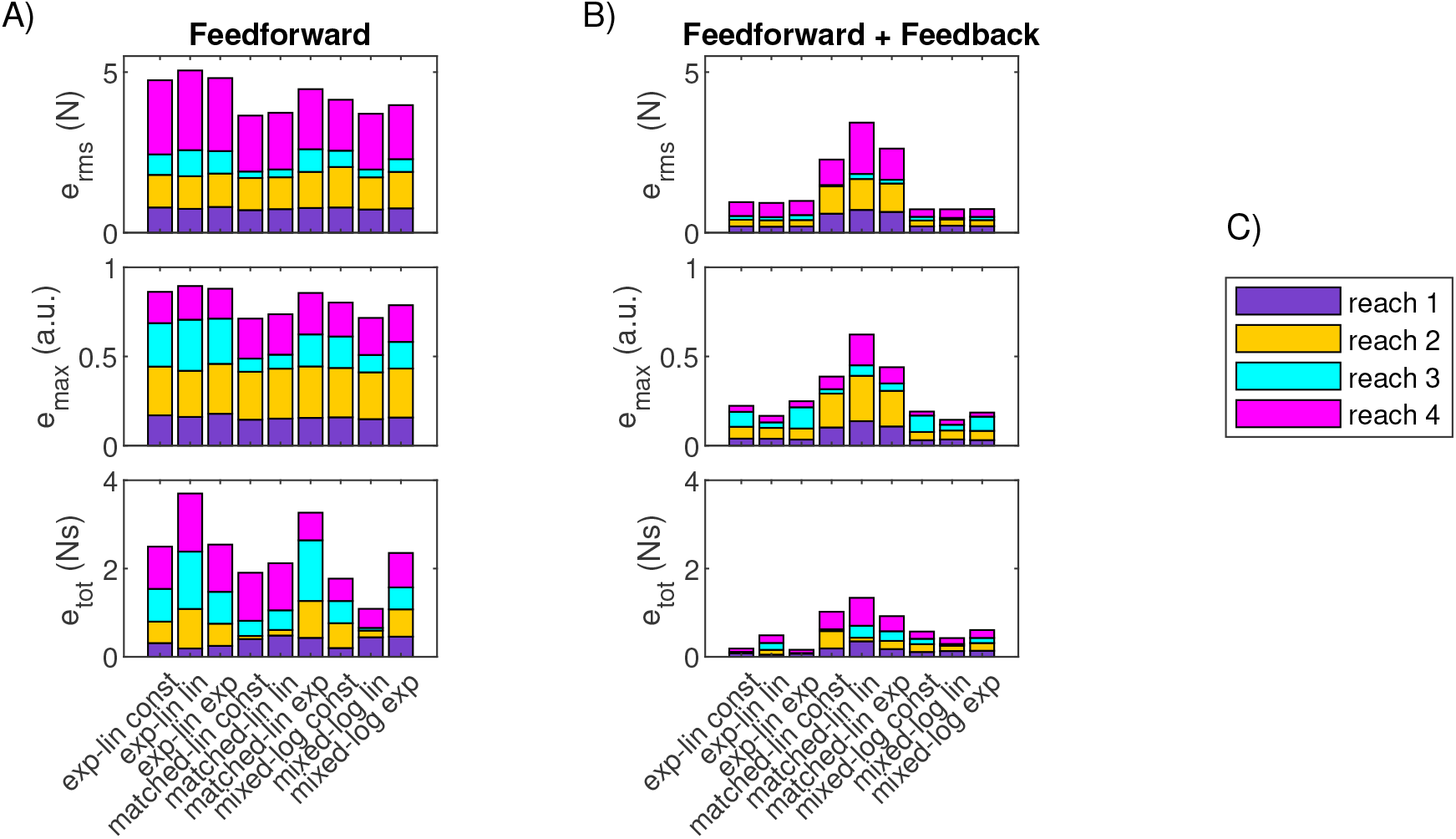
Force matching errors for the reaching tasks for feedforward and combined feedforward- feedback control. (A) Stacked performance errors for the nine MU pool models of the SF for the four reaching tasks under pure feedforward control. (B) Stacked performance errors for the nine MU pool models of the SF for the four reaching tasks under combined feedforward-feedback control. (C) Legend for task colours in panels A and B.

## 4 Discussion

The present work aimed to determine how MU pool structure and recruitment strategy affect the fidelity of force matching for various mechanical tasks. Towards this goal, we simulated the performance of nine different MU pool models in two different cases: an isometric trapezoidal force task for the TA and a simulated reaching task including predetermined length and contraction speed changes for the SF. In general, our simulations using a non-task-specific neural drive mapping and sequential MU recruitment produced force patterns which closely matched desired force profiles. Further improvements in the force match could be made by including a feedback component in the neural drive. These force matches arose from direct simulation of physiological phenomena (rather than curve fitting), providing evidence that this approach can be useful for future forward dynamics studies of neuromuscular control.

Our simulation results suggest that the structure and characteristics of MU pools, represented by the different models, can impact the pools’ ability to match a given force profile, but the differences are relatively small. We make five observations, in particular. 1) Relatively good force matches were obtained using a pure feedforward configuration, relying on a simple mapping from desired force to neural drive to govern MU recruitment and rate coding. 2) Enforcing onion-skin type behaviour through lower *r*_*max*_ of larger MUs had negligible impact on a MU pool’s overall force matching accuracy across the tasks studied. 3) Of the MU pools studied, matched-linear models performed the best in slowly varying isometric contractions under pure feedforward control, but this advantage was diminished when feedback was enabled. 4) The firing rate behaviour of exponential-linear pools during trapezoidal tasks exhibited the best qualitative match with experimental data. 5) There was very little overall difference between the pools’ performance across a range of non-isometric reaching tasks under pure feedforward control, but exponential-linear and mixed-logarithmic pools produced better force matches when feedback was enabled.

Functionally, one of the main differences between the MU pool models is the linearity of their steady-state response (Figure 4). Insight into the role of this linearity in force matching performance is provided by the matched-linear pool type, which was artificially constructed to produce linear steady-state input-output behaviour. Unsurprisingly, matched-linear pools performed better than the other pool models in the feedforward trapezoidal tasks requiring slowly varying force production. In contrast, steady-state linearity did not appear to have an advantage in reaching tasks, where the transient response of the MU pool dominates due to dynamically changing force demands; in fact, they exhibited worse performance relative to the other pool types when feedback was enabled. The physiologically motivated MU pool models (exponential-linear and mixed-logarithmic types) were more responsive to feedback but the root causes of this are challenging to study due to the dependence of MU pool transient responses on contractile history and the exact timing of events. However, one major factor that could account for this observation is the number of overlapping recruitment bands. When recruitment bands overlap, a small change in neural drive will trigger changes in the firing rates of the corresponding MUs, but the time it takes for each MU to respond with an earlier or delayed excitation impulse depends on when the MU was last fired. The physiologically motivated MU pool types have a higher number of overlapping recruitment bands than the matched-linear pool types, particularly at low force levels, which increases the likelihood that at least one MU can respond to a feedback signal in a relatively short time. The recruitment bandwidth overlap pattern of exponential-linear and mixed-logarithmic MU pools is, however, one of the factors underlying their nonlinear steady-state input-output behaviour, suggesting a potential trade-off between desirable steady-state and transient control characteristics.

One of the main limitations of the present study is that different MU pool models are difficult to compare due to uncertainties of the MU pool parameterisation and the coupling between some of the threshold and rate parameters. Our choice to use uniform first and last recruitment thresholds across the MU pool models ensures comparability of the range of thresholds. However, in the case of the TA, it forced most of the MUs in exponential-linear pools to operate at firing rates well below their *r*_*max,i*_ values during the trapezoidal tasks, leading to force fluctuations, while also causing MUs in matched-linear pools to reach maximum firing rates nearly instantaneously. Furthermore, some parameters, such as minimum and maximum firing rates, can have different roles in modelling and exprimental settings. Experimental maximum firing rates, for example, tend to be lower during voluntary contractions compared to those obtained with intraneural stimulation [46] and reported maximal rates can be exceeded during double discharges or ballistic movements [4, 8]. In contrast, *r*_*max,i*_ values in models are hard, task-independent limits on motor neuron function, and the minimum and maximum firing rates exhibited by MU pool models during a force production task depend on factors beyond these parameter values as illustrated by Figure 7. Hence, while parameterisation based on experimental data is always desirable, it is not always feasible; factors such as differences between muscles [10, 46], individual variation (e.g. [45]), and task-dependent behaviour [46] of MUs pose challenges on combining data from different experiments in a way that is relevant for the models considered. A detailed sensitivity analysis, while outside the scope of the present study, could provide further insight into the functional consequences of both modelling choices and biological variation. Given our goal of laying the foundations for using multi-MU muscles in MSK modelling, our results highlight the need to consider MU pool parameterisation as a whole from the perspective of functionality.

Variation in contractile properties of MU pools is outside the scope of the current investigation to limit the number of confounding, often difficult-to-parameterise variables. However, differences in slow- and fast-twitch MU properties could be integrated into future MU pool models through, for example, varying the twitch contraction time continuously with MU number [13] or using different distributions to draw the properties for the low-threshold slow-twitch MUs and the high-threshold fast-twitch MUs [17]. Differences in maximum contraction speed, i.e. the speed at which muscles fibres can no longer produce force due to the speed of shortening, could also be included, as has been done for two-MU models [31, 33], by specifying tasks in terms of muscle length rather than FVL trajectories. In our study, the impact of variation in contractile properties on the presented results would likely be small due the particular demands of our chosen tasks. Specifically, the low forces required for the reaching tasks result in only the smallest slow-twitch MUs being recruited whereas the isometric and slowly-varying forces in the trapezoidal tasks are dominated by tetanic rather than twitch characteristics. However, a more extensive cross-comparison between muscles and tasks or simulations of high-force non-isometric tasks would require addressing the potential functional impact of differences in contractile properties.

The mapping from desired force to neural drive is another feature of the present model that is difficult to relate to experimental data and hence subject to uncertainty. Although the form of Eq (2)) is appealing because of the minimal number of parameters, it only allows for a limited manipulation of the desired force and FVL signals to obtain the feedforward neural drive. The results of the parameter optimisation suggest that a mapping that cancels out nonlinearity of the each pools’ steady-state input-output behaviour (i.e. mapping from neural drive to tetanic muscle force) leads to best results, and furthermore, simulations with feedback enabled suggest that real-time feedback can successfully mitigate the effects of MU pool transient behaviour. However, this does not rule out the possibility that performance could be enhanced with a markedly more complex feedforward mapping, for example, in situations where feedback is less effective due long delays. Alternatively, the feedforward performance could be improved by obtaining multiple neural drive mappings focusing on narrower force ranges or single contraction types, although these run the risk of overfitting the simulation dynamics to the specific demands of the elementary tasks, or by explicitly mapping and inverting the steady-state nonlinearity (as done in [23]). These approaches are not, however, guaranteed to improve force matching in transient-dominant tasks, without reliance on feedback.

True MU behaviour is complex, and for the MU pool models studied, some known effects had to be omitted. Motor neuron firing is a stochastic process, and most MU pool models include noise in the timing of the excitation impulses following [13]. Motor signal noise was left out of the present study to simplify parameter optimisation and comparison of force matching results in tasks where force demand changes rapidly and small differences in the timing of individual impulses can lead to large changes in output. The investigated MU pools also treat recruitment and derecruitment identically, whereas current-voltage hysteresis exhibited by real motor neurons leads to different thresholds and firing rates when muscle force generation is increasing and decreasing [1, 48]. While this effect can be incorporated by utilising different motor neuron properties during the descending phase of trapezoidal tasks (as done by, e.g., [12]), it is less clear how practicable this approach is for force profiles with multiple inflection points, such as our reaching tasks, especially as repeated activation can affect the hysteresis observed [48]. The effects of including these and other MU characteristics, such as fatigue and other time-dependent changes in MU behaviour (see e.g. [18, 19, 20] for models including them), are left for future studies.

In the present study, force matching has been looked at in isolation from its natural dynamic context. This approach enabled comparing different MU pools under strictly comparable non-isometric conditions. However, until future studies embed MU pool models in a forward dynamics MSK model, the significance of the force matching errors is difficult to evaluate. In a full forward dynamics model, the effects of a force mismatch could be partially mitigated by the intrinsic stabilising properties of muscle FVL characteristics [49, 50, 51]. Furthermore, the presence of redundant and antagonistic muscles would reduce the demand on an individual muscle to produce the desired force. Rather, the focus of the task would shift to coordinating the recruitment of multiple muscles to match desired joint torque, which might allow individual MU pools to find favourable operating points. Our simulation results suggest that the addition of a small feedback element to a simple feedforward control scheme can improve force matching notably, and this would be expected to be generally true in MSK simulations as well. However, physiological motor control feedback loops have intrinsic sources of delays and noise, which were not modelled in our feedback scheme, and accounting for these factors might limit the amount of performance improvements feedback can deliver. Delays, in particular, can be a source of instability in feedback control [52], and whether a multi-body MSK system with antagonistic actuation would attenuate or amplify such instabilities during a complex movement is difficult to predict. Hence, incorporating MU pool models in forward dynamics simulations is the next step towards understanding the extent to which inherent MU pool characteristics affect movement functionality.

In conclusion, our investigation signals that muscles consisting of multiple rate-coded MUs can match forces from various mechanical tasks, building the foundation for incorporating these models in MSK modelling. These MU pool models offer a significant advance over the non-physiological amplitude-coded single-MU muscle models in terms their realism, and they also open a range of new research questions and topics for investigation, such as the correspondence between the natural diversity in MU numbers and pool characteristics [10] and differences in function across muscles. Using traditional approaches, evaluation of the functional differences among the muscles with diverse MU populations would not be feasible.

## Funding

This research was funded by the Wellcome Trust Investigator Award 215618/Z/19/Z.

## Supplementary Information

**Supplementary Figure 1. Figure 11: Supplementary Figure 1.**
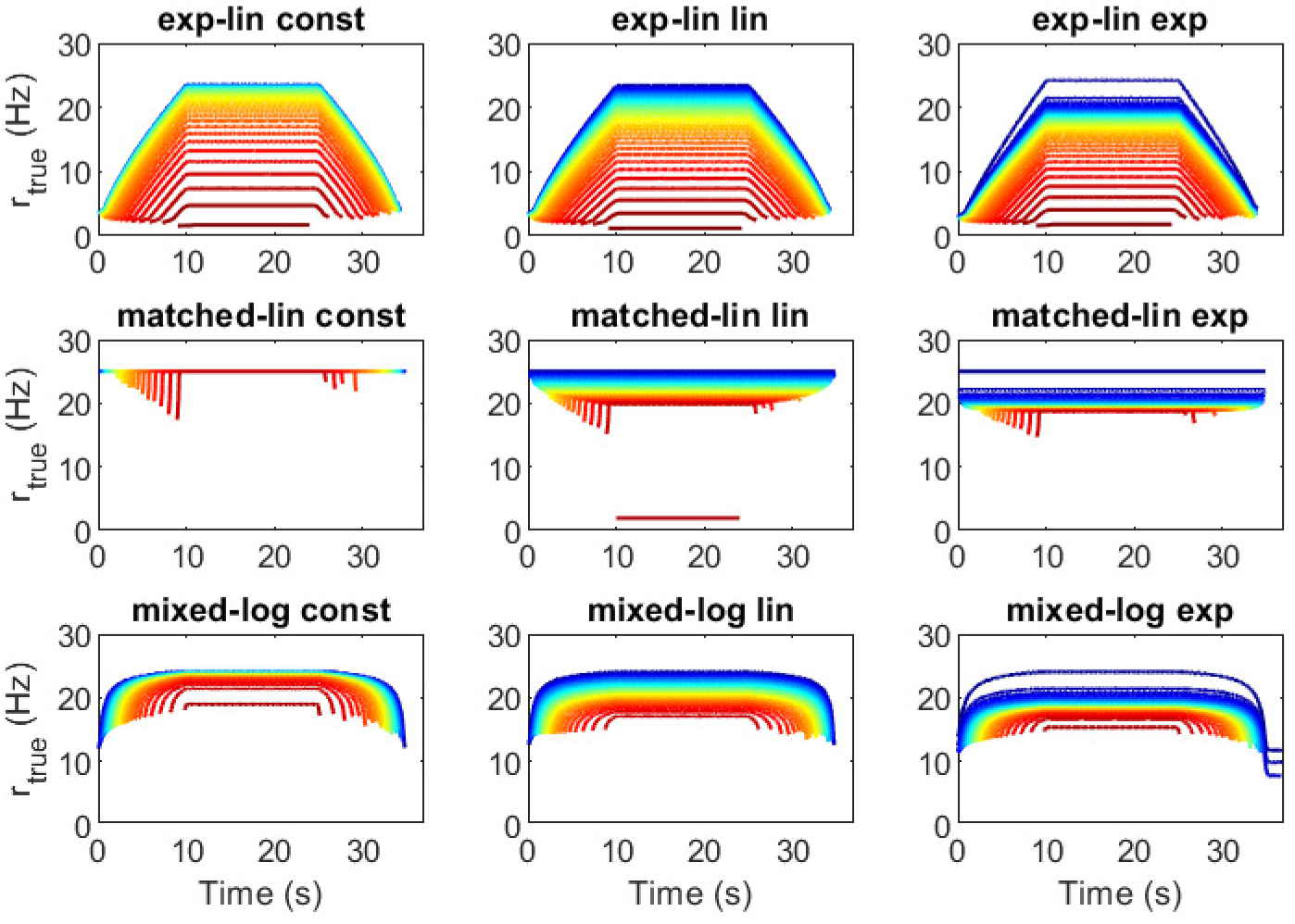
Firing frequencies for the TA/trapezoidal 0.5*F*_*max*_ task extracted from the excitation impulse trains for the nine MU pool models. The colours in all panels indicate order of MU recruitment from first (dark blue) to last (dark red). Only every 10^th^ MU is shown.

**Supplementary Figure 2. Figure 12: Supplementary Figure 2.**
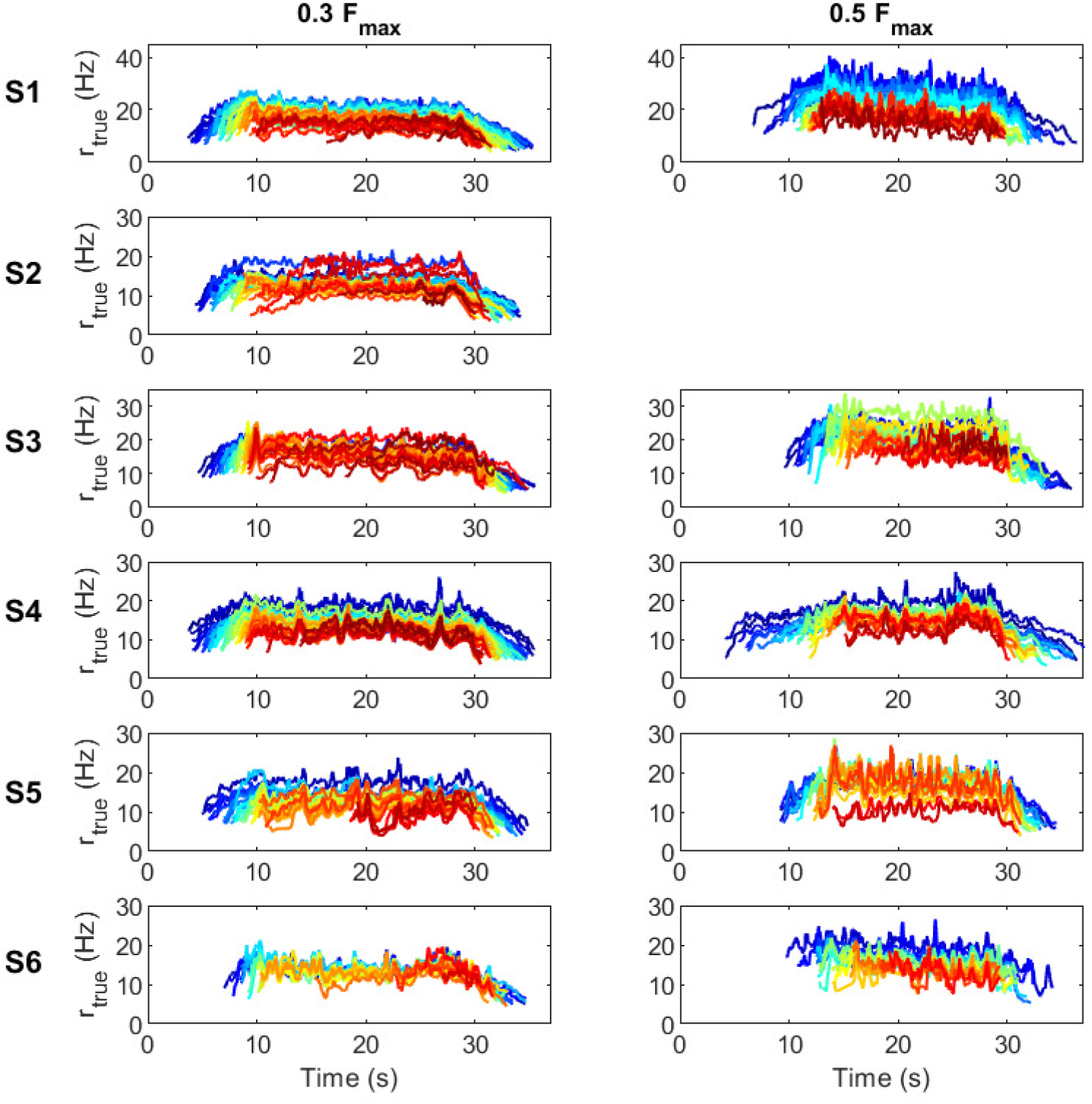
Experimental firing frequencies during trapezoidal task. Data on firing instants from [5] is converted into firing rate curves by computing inter-impulse distances and using a moving average filter. Both 0.3*F*_*max*_ (left column) and 0.5*F*_*max*_ (right column) tasks are shown for six subjects (S1-S6), except the higher force task for subject S2, which was not included in the original dataset.

## Notes

### Competing Interest Statement

The authors have declared no competing interest.

### Summary of Updates

Combined feedforward-feedback control added as control strategy with related updates throughout the manuscript. Methods rearranged for clarity. Calculation of discrete time correction term updated.

